# SpatialQPFs: An R package for deciphering cell-cell spatial relationship

**DOI:** 10.1101/2024.06.17.599458

**Authors:** Xiao Li

**Affiliations:** Computational Science and Informatics, Roche Diagnostics Solutions, Santa Clara, CA 95050, USA

**Keywords:** spatial statistics, cell-cell relationship, computational pathology

## Abstract

Understanding spatial dynamics within tissue microenvironments is crucial for deciphering cellular interactions and molecular signaling in living systems. These spatial characteristics govern cell distribution, extracellular matrix components, and signaling molecules, influencing local biochemical and biophysical conditions. Despite significant progress in analyzing digital pathology images, current methods for capturing spatial relationships are limited. They often rely on specific spatial features that only partially describe the complex spatial distributions of cells and are frequently tied to particular outcomes within predefined model frameworks. To address these limitations, we present *SpatialQPFs*, an R package designed to extract interpretable spatial features from cell imaging data using spatial statistical methodologies. Leveraging segmented cell information, our package offers a comprehensive toolkit for applying a range of spatial statistical methods within a stochastic process framework, including analyses of point process data, areal data, and geostatistical data. By decoupling feature extraction from specific outcome models, *SpatialQPFs* enables thorough large-scale spatial analyses applicable across diverse clinical and biological contexts. This approach enhances the depth and accuracy of spatial insights derived from tissue data, empowering researchers to conduct comprehensive spatial analyses efficiently and reproducibly. By providing a flexible and robust framework for spatial feature extraction, *SpatialQPFs* facilitates advanced spatial analyses, paving the way for new discoveries in tissue biology and pathology.

## Introduction

Understanding the spatial characteristics within tissue microenvironments is crucial for unraveling the intricate dynamics of cellular interactions and molecular signaling within living systems. These spatial patterns dictate the distribution of cells, extracellular matrix components, and signaling molecules, shaping the local biochemical and biophysical environment. Deciphering these spatial characteristics provides critical insights into various physiological and pathological processes, including tissue development, homeostasis, disease progression and ultimately associates with patient clinical outcomes^1–4^. By discerning spatial relationships between cell types, researchers are offered valuable insights into tissue architecture, cell-cell communication networks, and microenvironment dynamics, facilitating the identification of spatially defined biomarkers and therapeutic targets for precision medicine approaches^5–7^.

With the advance of imaging technology, digital pathology imaging of tumor tissue slides, such as those stained with Hematoxylin and Eosin (H&E) is becoming a routine clinical procedure for cancer management. This process produces massive information that capture histological details in high resolution. Thanks to the recent advances in artificial intelligence, automatic detection of different cell identities and corresponding spatial coordinates are made possible at large scale. Applying principles and advanced quantitative methods on these captured cell data, e.g. spatial statistics, can suggest novel solutions to fulfill this need. In recent years, there has been a significant interest in leveraging these cellular-level data to understand cell-cell spatial relationships and their association with various clinical outcomes, which have produced encouraging findings^5,8–21^. For example, Vu, T. et al.^20^ exemplifies this trend by utilizing an additive functional Cox regression model to analyze survival outcomes based on specific spatial features, such as Moran’s I and Mark connection function. Xiong, J. et al.^19^ developed an R package based on Ripley’s H function for interactive marker gating, which enhances the ability to identify and categorize cellular markers within spatial data. Canete, N. et al.^21^ presented spicyR, a tool for identifying differential cell-type colocalizations across different groups by employing Ripley’s L function, thereby enhancing the understanding of how cellular environments change with disease pathology. Seal, S. et al.^17^ leveraged point process models (Ripley’s K, L function, and Pair correlation function) to quantify spatial co-occurrence and build Functional Analysis of Variance (ANOVA) model for statistical inferences on spatial relationships between cell types across distinct disease groups.

While these previous efforts in spatial analysis of cell-level data have provided valuable insights, they are limited by their reliance on specific choices of spatial features. The use of features derived from spatial point process models versus spatial lattice process models can significantly influence the characterization of spatial properties between cell objects. Spatial point process features are advantageous in capturing fine-resolution spatial behaviors, as they model the precise locations of cells within a tissue region, allowing for a detailed understanding of spatial interactions and distributions. Conversely, spatial lattice process features, which treat the data as areal or aggregated, offer a broader, more generalized view of spatial relationships. While this approach is useful for understanding large-scale patterns, it may limit the ability to detect and characterize finer-scale spatial dynamics. Additionally, these efforts often overlook geostatistical features, which are essential for capturing continuous spatial variation and understanding spatial dependency or correlation across distances. This omission restricts the potential for a more comprehensive analysis of spatial heterogeneity and multiscale interactions within the tissue microenvironment. Moreover, these approaches are often tied to particular outcome models, such as regression models for continuous outcomes or survival models for time-to-event outcomes. Consequently, they tend to focus on narrow applications, thereby limiting the broader clinical applicability of spatial analysis.

To overcome these limitations, we present *SpatialQPFs* (**S**patial **Q**uantitative **P**athology **F**eature**s**), a comprehensive and extendable R package designed for deciphering spatial characteristics within and between cell types in the tissue microenvironment. By leveraging advanced imaging analysis techniques like tumor region-of-interest (ROI) identification, cell segmentation and classification, it becomes feasible to convert the raw, information-rich digital pathology images into a structured tabular format containing cell-level details, such as cell coordinates and identities. *SpatialQPFs* then, from a holistic point of view, take this cell level data as input and organizes it as point process data, areal data and geostatistical data, respectively, where a collection of spatial statistics methodology can be employed to extract a broader array of human interpretable quantitative pathology features, empowering researchers to conduct comprehensive spatial analyses for deciphering cell-cell spatial relationship efficiently and reproducibly. Furthermomre, by decoupling the extraction of spatial features from specific outcome models, *SpatialQPFs* empowers researchers to explore diverse clinical and biological questions without being restricted to predefined frameworks, thus expanding the potential for spatial analysis in various disease contexts and research applications. This flexibility and comprehensiveness position our approach as a valuable tool for a wide range of implementations, including supervised, unsupervised, and semi-supervised tasks in computational pathology.

### SpatialQPFs R Package

The *SpatialQPFs* package is written in R^22^, an open-source, functional language and environment designed for statistical exploration and data visualization. Users can develop and customize the provided functions in the *SpatialQPFs* source code for various purposes and use cases.

The package primarily focuses on examining cell-cell spatial relationships. However, its functions are also relevant in diverse clinical contexts, including but not limited to analyzing spatial patterns of brain lesions^23,24^.

The major functions of the *SpatialQPFs* package, detailed in Table 1, are geared towards streamlining the analysis of structured tabular data at the single-cell level. This is achieved through the organization of data into various spatial data types, i.e. point process data, areal data and geostatistical data, each accompanied by visualization options that enhance data interpretation. For guidance on how to use these functions, refer to the section “Example workflow” below.

**Table 1.** Primary SapatialQPFs functions (see package manual for further details).

| Function | Description |
| --- | --- |
| Data_Vis() | Visualizes the input cell level data, including raw spatial map of individual cells and smoothed spatial density map |
| Point_pattern_data_uni() | Main function to generate spatial features using point process data (single cell type). |
| Point_pattern_data_bi() | Main function to generate spatial features using point process data (pairwise cell types). |
| Point_pattern_data_ITLR() | This is a function that discriminates a target cell population into 2 subgroups, in spatial relationships with the reference cell population. In case of lymphocytes and tumor cells, the lymphocytes are discriminated into intra-tumoral lymphocytes and adjacent-tumoral lymphocytes. |
| Areal_data() | Main function to generate spatial features using areal data. |
| Geostatistics_data() | Main function to generate spatial features using geostatistical data. |

### Data preparation

Digital pathology image analysis begins with several image preprocessing steps to ensure that the data is clean and accurately represents the biological structures of interest. These steps include quality control measures such as artifact removal, identification of tumor regions of interest (ROIs), and performing cell classification and segmentation. Once image processing is complete, the resulting cell mask from the cell classification and segmentation step is converted into a structured tabular format, which includes x- and y-coordinates along with the identified cell types. This tabular data provides a comprehensive and organized representation of the spatial distribution and characteristics of the cells within the image.

With the structured cell data prepared, *SpatialQPFs* proceeds to perform spatial feature extraction and visualization by leveraging this cell-level information to analyze the spatial relationships and patterns among the cells. To visualize the cell-level data and its derived spatial density map, researchers can use the`Data_Vis()` function. This visualization tool allows researchers to explore and interpret the spatial organization of the cells, facilitating a deeper understanding of the spatial layout and interactions within the tissue microenvironment.

In the following paragraphs, we will introduce how the three spatial data types, i.e. point process data, areal data, and geostatistical data, can be leveraged based on the structured cell-level data. We will also discuss the types of methods that can be employed for each spatial data type.

### Point process data

Spatial point process analysis is a vital technique in spatial statistics, used to examine the distribution of point objects such as cell nuclei and their spatial interplay. This method is crucial for understanding how cells are organized within a given tissue, which can provide insights into various biological processes and disease mechanisms.

The raw single-cell level input can be directly used for point process data analysis. To leverage point process methodologies in characterizing the spatial distribution of a singular cell type, *SpatialQPFs* provides the `Point_pattern_data_uni()` function. This function employs a series of techniques such as Ripley’s K-function, G-function, and Clark and Evans nearest neighbor analysis to determine whether the distribution of the specified cell type is random, clustered, or uniform^25,26^ in the tissue microenvironment. These analyses help in identifying patterns within the spatial arrangement of cells, which can be critical for interpreting biological phenomena. Furthermore, to quantify the spatial interplay between two cell identities, such as lymphocytes and tumor cells, *SpatialQPFs* includes the `Point_pattern_data_bi()` and `Point_pattern_data_ITLR()` functions. These functions utilize a variety of spatial point process methods, including the multi-type Ripley’s K-function, differences in Ripley’s K-function, multi-type G-function, Pair correlation function, Mark correlation function, Mark connection function, Marcon and Puech’s M function, and the cell population discrimination analysis. These methods provide detailed insights into the spatial relationships between different cell populations within the tissue, revealing interactions that may be crucial for understanding tumor biology and immune response^10,11,25,27–29^. Implementation details of these functions are provided in the “Example Workflow” section below, and the features derived from these point process methods, along with their interpretations, are detailed in Supplementary Material Section 1.

### Areal data

To apply spatial statistics methods for areal data effectively, a conversion process from the raw single-cell level input is required, involving the partitioning of the study area into a grid of subregions or lattices, each annotated with its corresponding geographic location, typically denoted by the coordinates of the grid centroid.

*SpatialQPFs* facilitates this process through its `Areal_data()` function, which partitions the spatial domain into square grids behind the scenes. Users can customize the grid size using the `scale` argument, achieving a balance between resolution for detecting local-regional variations and the stability of estimates. Guidelines for optimizing grid size have been previously reported in studies such as Page, D. B. et al.^30^, Mi, H. et al.^31^, Francis, K. et al.^32^, Turkki, R. et al.^33^ and Verghese, G. et al.^34^. Within each lattice grid, measurements related to the counts or percentages of each cell type are computed independently, resulting in a mosaic of estimations covering the entire tissue space. Autocorrelation analyses are then conducted to assess the degree of similarity between neighboring locations within the tissue sample for each distinct cell type. These metrics include global and local Moran’s I, Geary’s C, Getis-Ord statistics, Lee’s L statistics and their derived measures^9,35–39^. Such assessments help identify spatial clustering or dispersion patterns, revealing significant hotspots or coldspots in the distribution of cell types within the tissue, aiding in the identification of regions of elevated or reduced biological activity. Additionally, the degree of colocalization between different cell types is evaluated using methods such as the Morisita-Horn index, Sorensen index, and Jaccard index^40–42^. Implementation details of these functions are provided in the “Example Workflow” section below. Features derived from all the aforementioned methods and their explanations are detailed in Supplementary material Section 1.

### Geostatistical data

Geostatistics, a discipline comprised of mathematical and statistical methods, was originally devised to forecast probability distributions of spatial stochastic processes for mining operations. However, its utility has since expanded across a wide range of disciplines. It is utilized in the fields such as petroleum geology, earth and atmospheric sciences, agriculture, soil science, and environmental exposure assessment^16,43–49^. In recent years, geostatistical methods have also been employed in digital pathology images^50,51^. For instance, in a study on breast carcinoma, researchers introduced a novel measurement of heterogeneity using geostatistical methods for histopathological grading tasks^52^.

Geostatistical methods typically require continuous variables measured over space, making the direct application of these techniques to categorical data, such as cell types, challenging. To address this, *SpatialQPFs* adopted two strategies for applying geostatistical methods to cell-level data: 1. Create an indicator variable for a specific cell type, assigning a value of 1 if a cell belongs to that type and 0 otherwise. This binary variable can then be used to compute indicator variograms for indicator kriging^53^; 2. Transform the cell-level data into cell prevalence within each square lattice grid, similar to areal data analysis. By aggregating the data, one obtain continuous variables representing the proportion of cell types within each lattice grid. Semi-variograms can then be computed for ordinary kriging of each cell type, or cross-variograms can be used for co-kriging when analyzing the spatial relationships between two specified cell types^53^.

To employ geostatistics techniques for assessing the spatial characteristics of cells, the `Geostatistics_data()` function can be utilized to extract features that delineates the spatial variability for cell type(s) in the tissue microenvironment. The function integrates the Matérn variogram model^53^ to fit a theoretical variogram after computing the empirical variogram, which measures the variability between pairs of cells separated by different distances. This process unfolds in two key steps. Initially, the empirical variogram is computed, representing a discrete function that captures the variability measure between pairs of cells at varying distances. Subsequently, a theoretical variogram is fitted based on the estimated empirical variogram. From this fitted model, parameters characterizing the spatial variability of cells can be derived. Detailed implementation instructions for these functions are provided in the “Example Workflow” section below. Details regarding the geostatistics features extracted and their descriptions are provided in Supplementary material Section 1.

## Example workflow

In this section, we provide brief descriptions of the functionalities offered by *SpatialQPFs*. We will use the same H&E slide as an example throughout this section.

For first-time users, installation of *SpatialQPFs* can be completed in the user’s local computing environment:

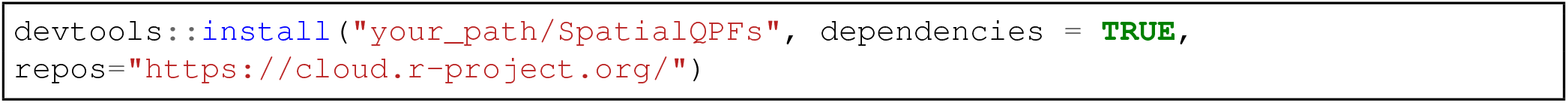

Then, the package can be loaded:

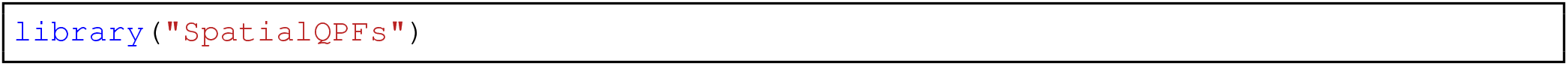

The utilization of the *SpatialQPFs* functionality begins by importing cell-level input data (currently only support.*csv* file, more input format will be supported in the future release). Preliminary steps, including identifying tumor regions of interest (ROIs), segmenting and classifying cells, are performed before invoking *SpatialQPFs* (Figure 1).

**Figure 1.**
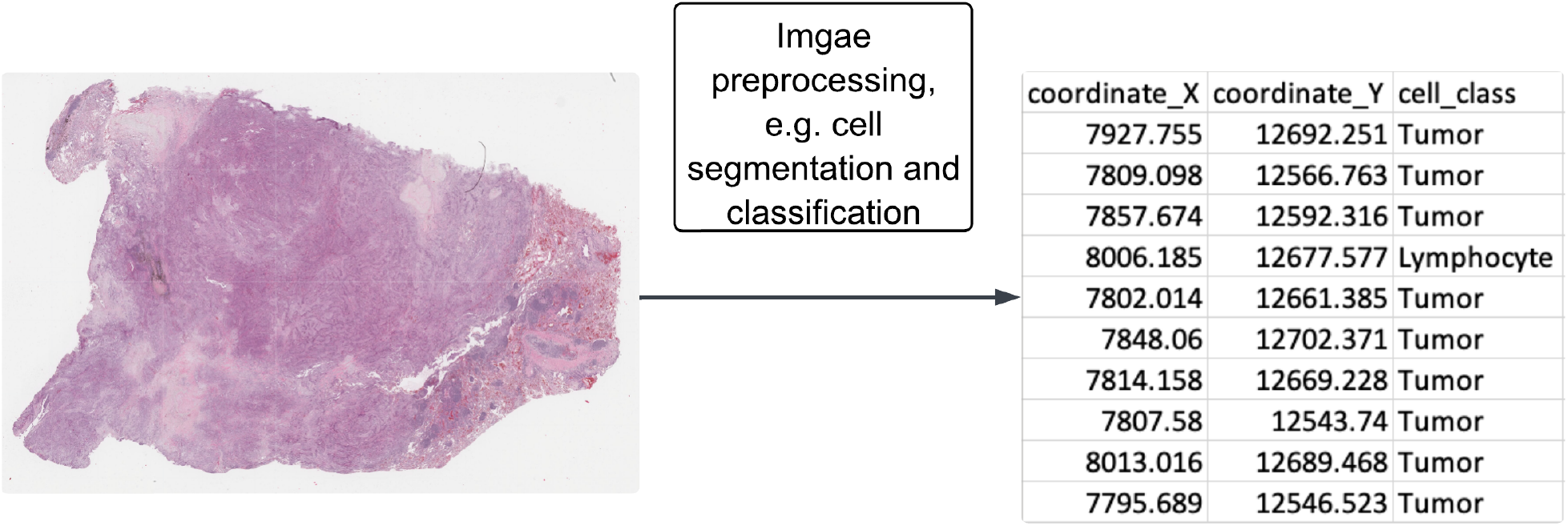
*SpatialQPFs* takes structured tabular cell-level information, such as the centroid coordinates of cells and their cell types, as input after image preprocessing steps are performed.

To visualize the spatial distribution of the input tabular data within the original tissue space and its derived spatial density map, users can call `Data_Vis()` function. For example, by specifying `cell_type =` “`Lymphocyte`” users can plot the lymphocyte population (Figure 2). The argument `path` and `file` accept strings that indicate the directory address where the file is saved and the name of the CSV file. Additionally, `cell_class_var`, `x_var` and `y_var` specify the column names from the input CSV file that contain the X, Y-coordinates of cell centroids and the corresponding cell type. See function manual for further details.

**Figure 2.**
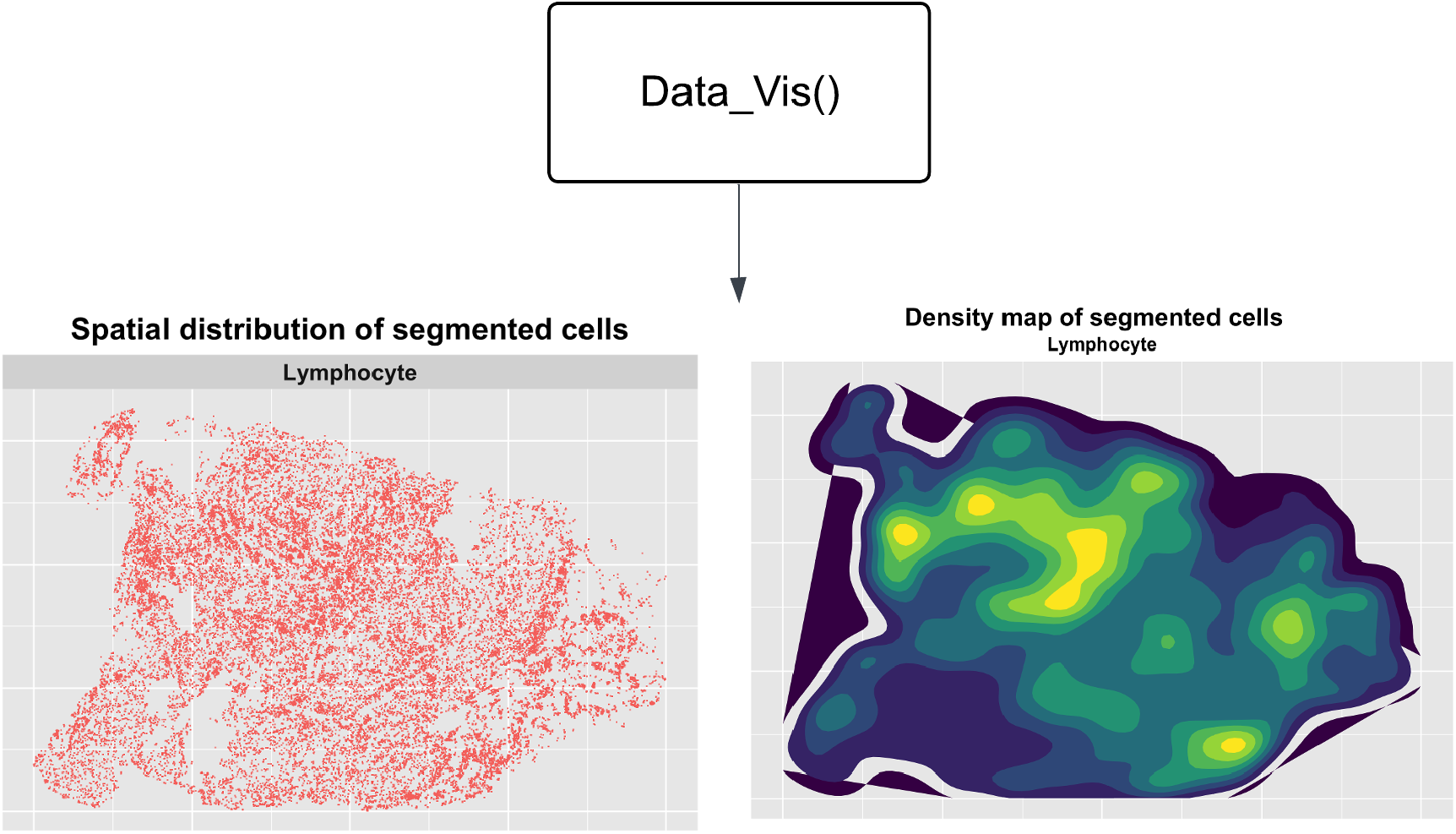
Spatial distribution of user specified cell type and its derived spatial density map.

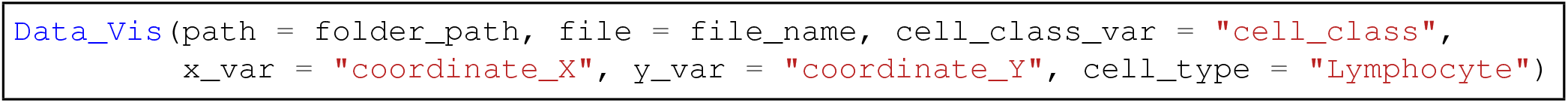

The analysis of such structured cellular-level data often begins with point process analysis. Users can start by calling `Point_pattern_data_uni()` to investigate whether the spatial clustering pattern of a single cell type is random, clustered, or uniform, over a range of distances, e.g. Ripley’s K Function. `scale` specifies the maximum distance, and the function internally generates a sequence of numbers from 0 to maximum distance, evenly spaced with a total of 10 values at which the point process data analysis are conducted. Further, spatial interplay of a pair of cell types can be analyzed by calling `Point_pattern_data_bi()`, which evaluates how the two cell populations are distributed relative to each other, e.g. Multi-type G Function. As the multiple type point process data analysis methods describes directional spatial relationship, `from_type` specifies the source cell type (e.g., Lymphocyte) and `to_type` specifies the target cell type (e.g., Tumor). Lastly, `Point_pattern_data_ITLR()` discriminates the source cell population into subgroups based on their spatial relationships with the target cell population. To illustrate, we use “Tumor” as the target cell population and “Lymphocyte” as the source cell population, the function then internally performs the following steps: 1) it profiles the global spatial distribution of tumor cells by quantifying their density using a kernel estimate, which builds a tumor landscape where ‘hills’ represent regions densely populated with tumor cells, such as tumor nests. The height of each hill correlates with tumor density at that specific location; 2) for each lymphocyte, its spatial proximity to tumor cells is quantified by referencing the tumor density landscape at its location. This provides an efficient quantitative measurement of proximity to tumor cells for every lymphocyte; 3) a two-component Gaussian mixture model (GMM)^54^ is applied to these measurements, discriminating lymphocytes into two distinct groups: ‘ITL’ (intra-tumor lymphocytes, where the associated tumor density is high) and ‘ATL’ (adjacent-tumor lymphocytes, where the associated tumor density is low). Then features like *n*_*ITL*_*/*(*n*_*ATL*_ + *n*_*ITL*_) or *n*_*ITL*_*/n*_*TC*_ which represent the proportion of intra-tumor lymphocytes relative to the total number of lymphocytes or the total number of tumor cells, can be calculated to assess the degree of colocalization between the two cell types.

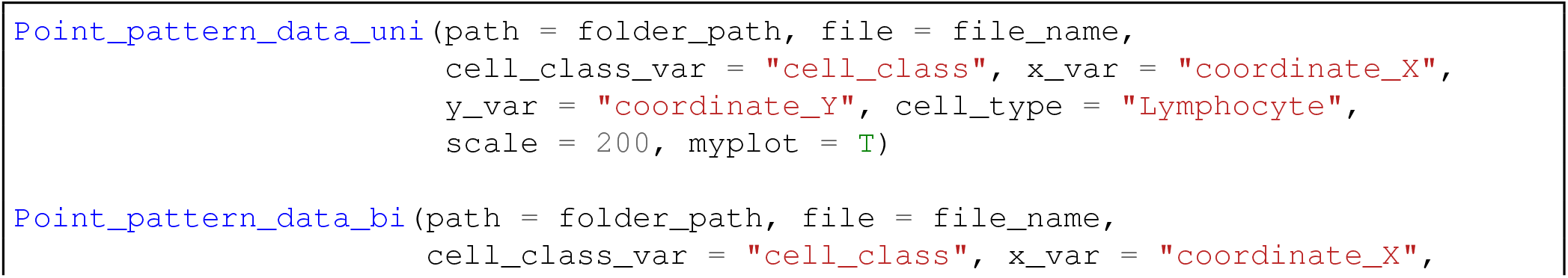

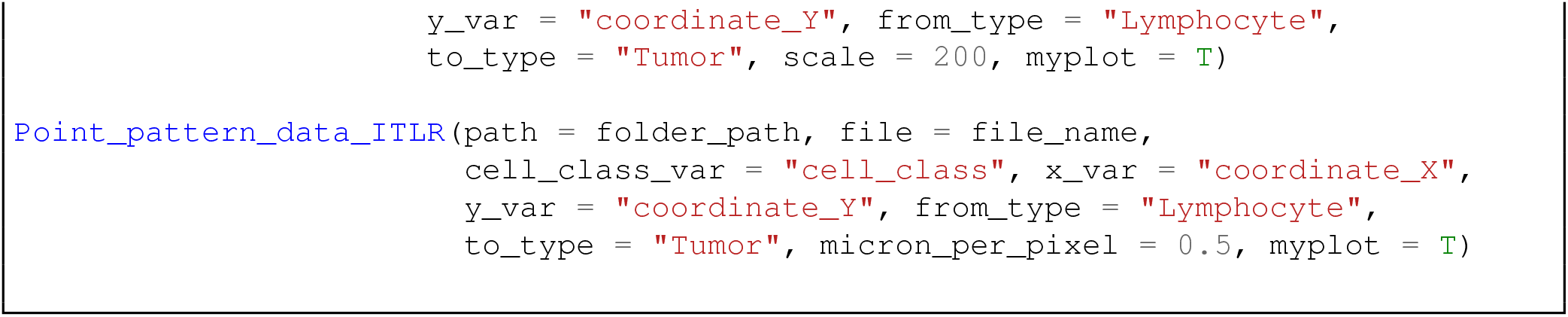

Figure 3 shows selected plots from point process analysis, including Multi-type G Function (top left) of Lymphocyte and Tumor cells, Ripley’s K Function for Tumor cells (top right) and the localization of Lymphocyte subpopulations (bottom).

**Figure 3.**
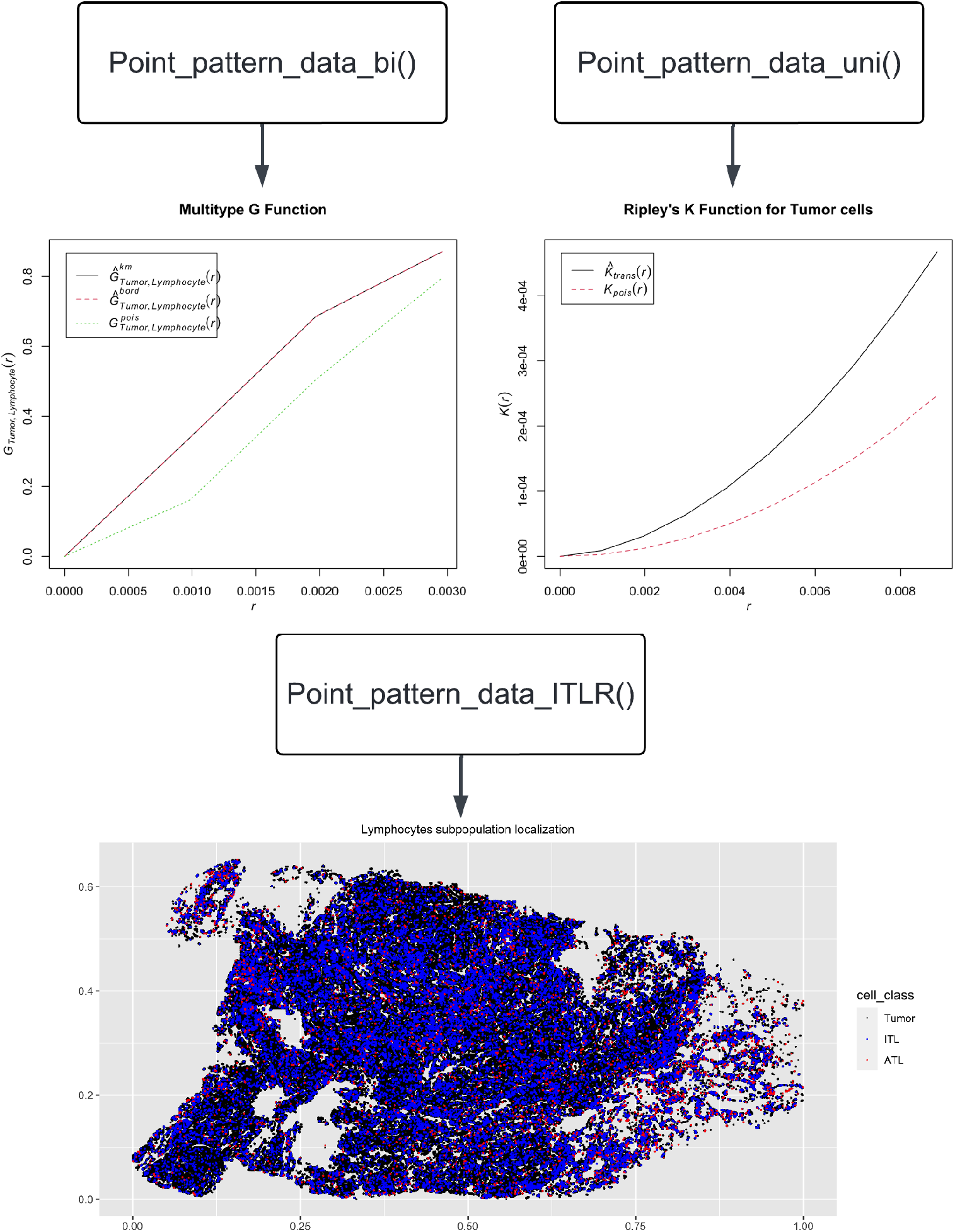
Selected plots from point process analysis, including Multi-type G Function (top left) of Lymphocyte and Tumor cells, Ripley’s K Function for Tumor cells (top right) and the localization of Lymphocyte subpopulations (bottom).

Additional plots for other point process data analysis methods are presented at https://github.com/Genentech/SpatialQPFs/blob/main/Tutorial-of-SpatialQPFs.html. Additional details of function arguments can be found in the function manuals.

Furthermore, the analysis of areal data can be performed using the `Areal_data()` function. Note that, this function conducts the transformation from point process data to areal data behind the scenes, by partitioning the underlying space into square lattices. Users can specify the desired side length of the square lattices as 2*`scale`.

Figure 4 (top) presents the prevalence map for tumor cells, with the tumor cell prevalence displayed at the center of each lattice grid. Higher values indicate regions with a denser tumor distribution. To identify local clustering pattern, various areal data analysis methods can be employed, e.g. local Moran’s I, to compute local measures of spatial autocorrelation for tumor cells (Figure 4 (bottom)), wherein the spatial lattices are categorized into “High-High”, “High-Low”, “Low-High”, “Low-Low”, and “non-significant” groups. This categorization is based on statistical inferences from Monte Carlo simulations, comparing the prevalence of tumor cells within each lattice to those in neighboring lattices. The percentage of each local pattern type can then be calculated as features. Similarly, features can be calculated from an analogical manner using local Geary’s C, Getis-Ord and Lee’s L statistics.

**Figure 4.**
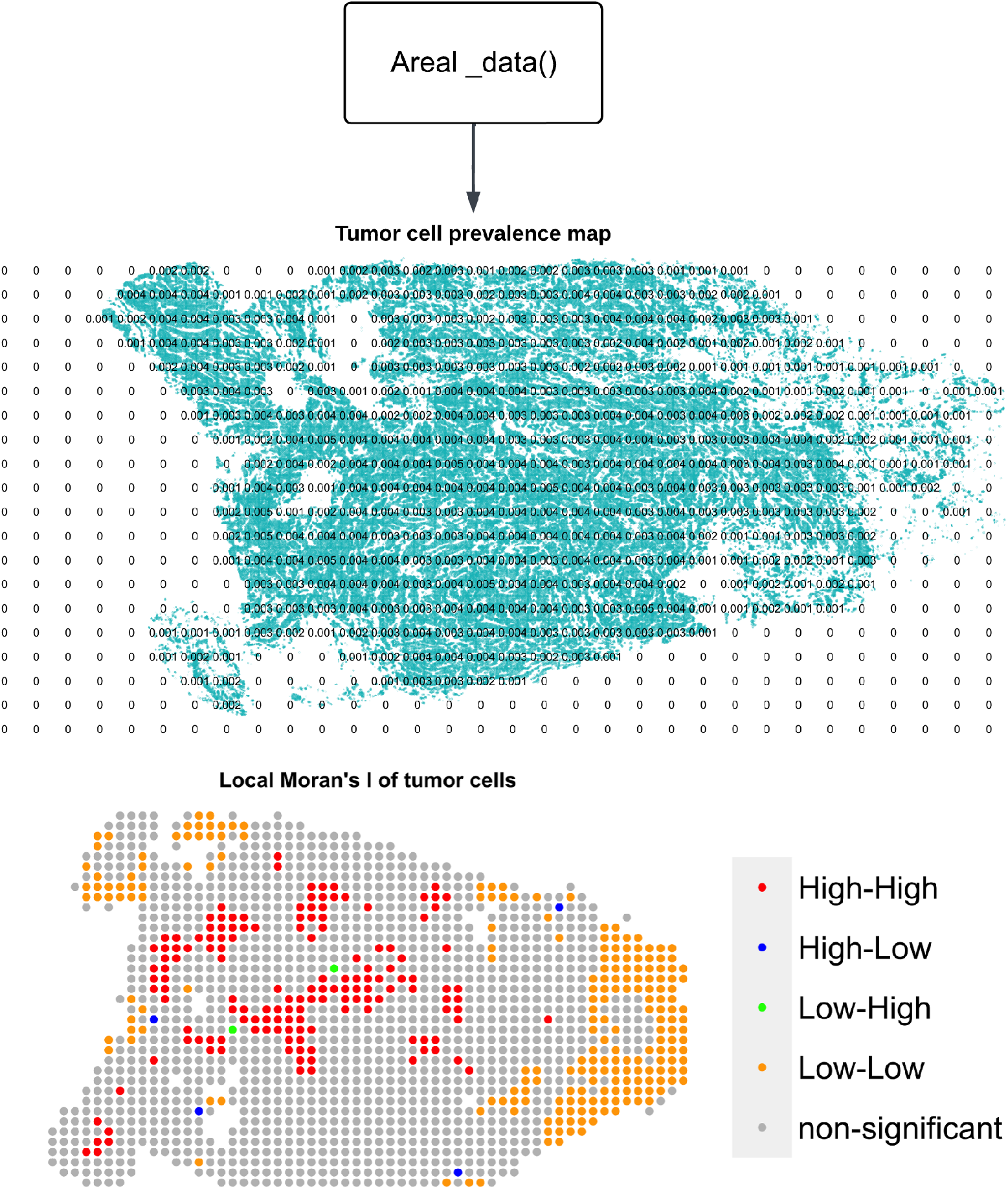
Tumor cell prevalence map and local Moran’s I for tumor cells, the spatial lattices are grouped to “High-High”, “High-Low”, “Low-High”, “Low-Low” and “non-significant” according to the prevalence of tumor cells within them and those of neighboring lattices.

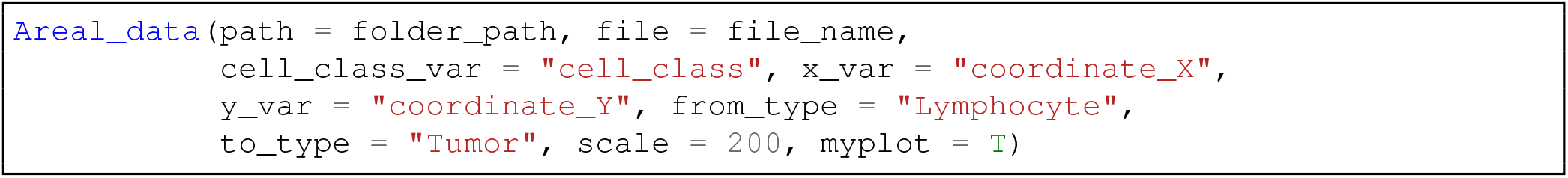

Other areal data analysis plots can be found in https://github.com/Genentech/SpatialQPFs/blob/main/Tutorial-of-SpatialQPFs.html. Detailed instructions for function arguments are available in the function manuals.

Finally, the analysis of geostatistical data can be conducted using the `Geostatistics_data()` function. In the current version of *SpatialQPFs*, indicator variogram^53^, semi-variogram^53^and cross-variogram^53^ are implemented to illustrate the spatial correlation of either a single cell type or a pair of different cell types, respectively. Specifically, to calculate the indicator variogram, as mentioned in the “Geostatistical Data” section, *SpatialQPFs* internally creates an indicator variable for a given cell type, either `from_type` or `to_type`, since the binary indicator variable applies identically to both cell types. Additionally, after transforming the cell-level data into cell prevalence within each square lattice grid, the semi-variogram can be calculated separately for `from_type` and `to_type`, while the cross-variogram is computed using both `from_type` and `to_type` together. Similar to areal data analysis, users can specify the side length of each grid as 2*`scale`, allowing flexibility in the spatial resolution of the analysis.

As depicted in Figure 5, a semi-variogram analysis of tumor cells is performed. Firstly, the empirical semi-variogram is calculated for tumor cell pairs that are separated by various distances. Following this, a smooth curve is overlaid to represent the fitted model that captures the spatial correlation behavior. The function internally employs Matérn variogram model for this analysis.

**Figure 5.**
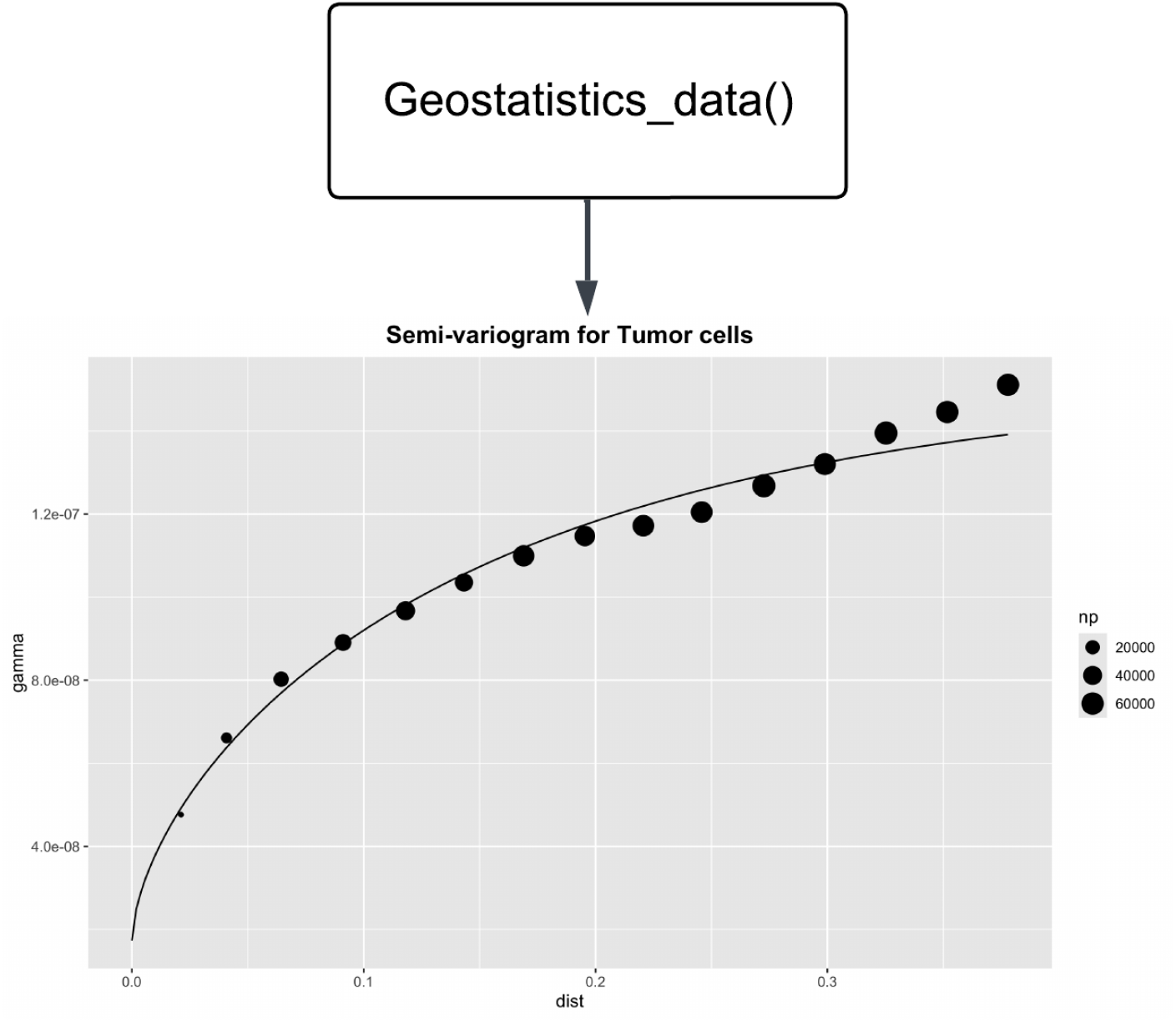
Semi-variogram analysis of tumor cells.

**Figure 6.**
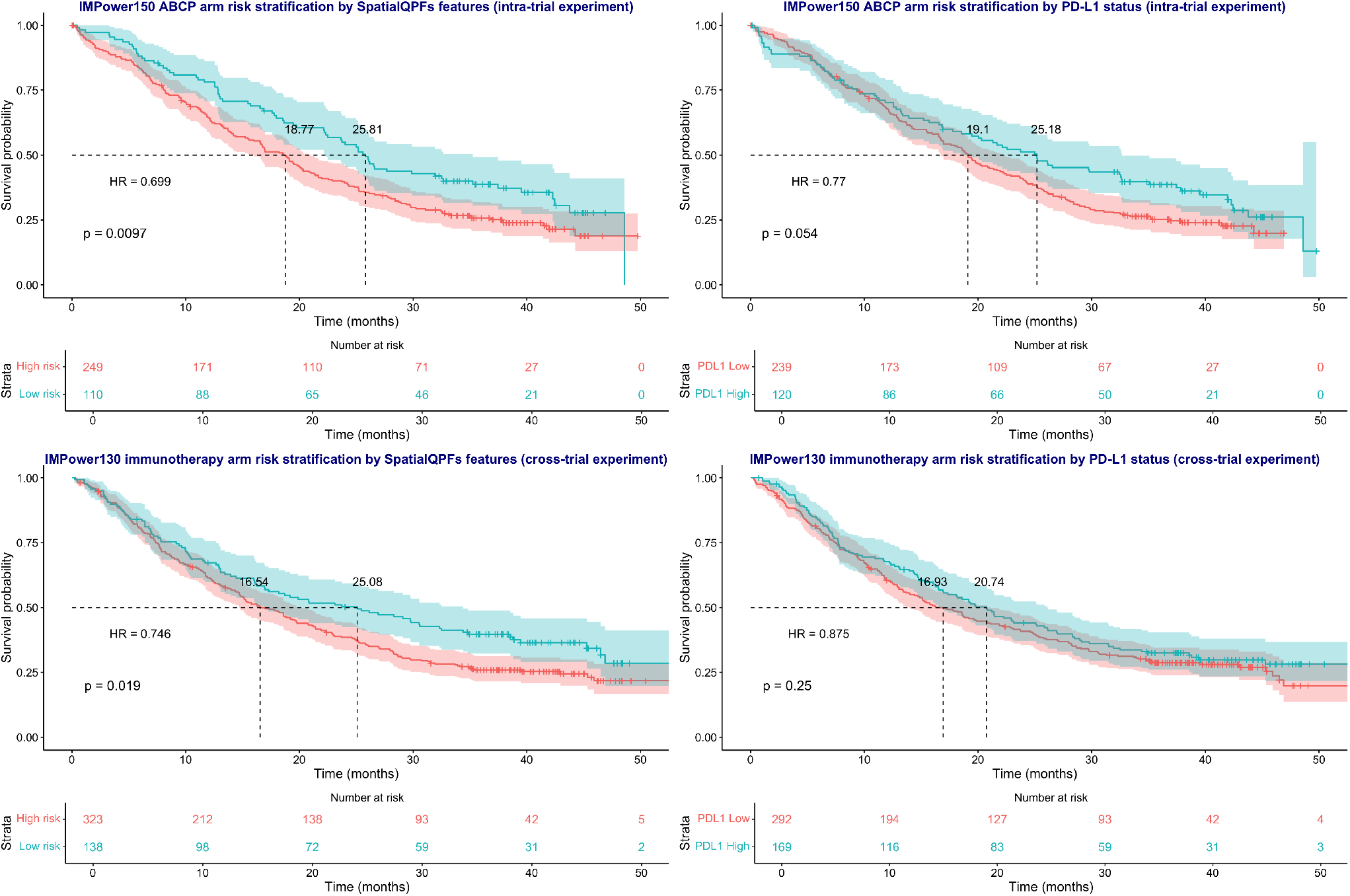
Kaplan-Meier survival curves illustrating risk stratification using SpatialQPFs features and PD-L1 status across two clinical trials: IMpower150 (intra-trial setting, top panels) and IMpower130 (cross-trial setting, bottom panels).

Further plots for alternative geostatistical data analysis methods are demonstrated at https://github.com/Genentech/SpatialQPFs/blob/main/Tutorial-of-SpatialQPFs.html. For detailed guidance on function arguments, refer to the function manuals.

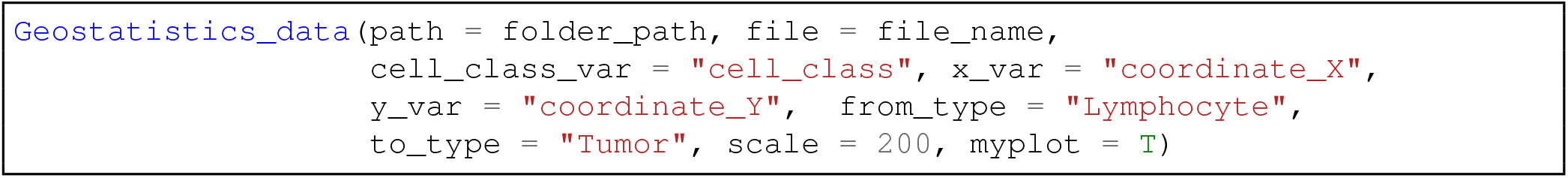

In Supplementary material Section 2, we present the additional R output that includes the full list of the features obtained from this example, by calling the functions `Point_pattern_data_uni()`, `Point_pattern_data_bi()`, `Point_pattern_data_ITLR()`, `Areal_data()`, `Geostatistics_data()`. The complete list of features, along with their meanings, is presented in Supplementary Material Section 1, reflecting the associated spatial methods employed.

### Applying SpatialQPFs to predict clinical benefit for cancer immunotherapy

To showcase the practical utility of *SpatialQPFs*, we utilized its capabilities to extract spatial features deciphering the spatial relationship between tumor cells and lymphocytes from all the available H&E stained digital pathology slides. These slides were sourced from two cancer immunotherapy (atezolizumab) clinical trials, IMPower150 (NCT02366143) and IMPower130 (NCT02367781), sponsored by F. Hoffmann–La Roche/Genentech (for details, refer to Supplementary material Section 3). Subsequently, we explored the association between these features and patients’ overall survival.

We designated the ACP arm (atezolizumab plus carboplatin plus paclitaxel, n = 381) of IMPower150 as our training set and the ABCP arm (atezolizumab plus carboplatin plus paclitaxel plus bevacizumab, n = 359) of IMPower150 as test cohort 1, for the intra-trial experiment. Additionally, we conducted a cross-trial experiment, utilizing the immunotherapy arm (atezolizumab plus Carboplatin plus Nab-Paclitaxel, n = 461) from IMPower130 as test cohort 2.

To construct the prediction model incorporating relevant spatial features, we first applied a univariate Cox proportional-hazards model^55^ to each of the features generated by *SpatialQPFs* (see the full list in Supplementary Material, Section 2) within the training set. We then performed a feature selection process to identify the most relevant predictors, using a relaxed p-value threshold of *p* ≤ 0.15. This threshold ensures that potentially important variables are not excluded prematurely, while maintaining an appropriate level of rigor. This approach is particularly useful during the exploratory stages of model building or when dealing with high-dimensional data, as recommended in previous studies^56,57^. To address collinearity among the selected features, we then applied L2-regularized Cox proportional-hazards regression^58^, a common technique to penalize model coefficients and mitigate multicollinearity. This approach allowed us to construct a robust prediction model using the selected spatial features to predict patients’ survival risks.”

The results of the intra-trial experiment (IMPower150 ABCP arm) demonstrated that the high-risk group, as stratified by *SpatialQPFs* features, exhibited significantly poorer overall survival compared to the low-risk group (HR = 0.699, logrank p = 0.0097). The median survival times were 18.77 months for the high-risk group and 25.81 months for the low-risk group. This indicates a strong association between the spatial features extracted by SpatialQPFs and patient outcomes in the ABCP arm of IMPower150. In contrast, stratification by PD-L1 status in the same arm (IMPower150 ABCP) showed a less pronounced difference in survival outcomes, with an HR of 0.77 and a logrank p-value of 0.054, suggesting a borderline significant trend but not a strong enough distinction between the PD-L1 high and low groups. The median survival times were 19.1 months for the PD-L1 low group and 25.18 months for the PD-L1 high group.

For the cross-trial experiment using the IMPower130 immunotherapy arm, the risk stratification by *SpatialQPFs* features again revealed a significant difference in overall survival between the high- and low-risk groups (HR = 0.746, logrank p = 0.019). The median survival times for the high-risk group were 16.54 months, compared to 25.08 months for the low-risk group, demonstrating the utility of *SpatialQPFs* in predicting patient survival across different clinical trials. However, similar to the intra-trial results, stratification by PD-L1 status in the IMPower130 immunotherapy arm yielded a weaker distinction, with an HR of 0.875 and a non-significant logrank p-value of 0.25. The median survival times were 16.93 months for the PD-L1 low group and 20.74 months for the PD-L1 high group, suggesting that *SpatialQPFs* may offer more robust risk stratification compared to PD-L1 status alone across both trials.

Overall, these results highlight the ability of *SpatialQPFs* to decipher tumor-immune spatial relationships, offering meaningful risk stratification for cancer immunotherapy that extends beyond traditional biomarkers such as PD-L1.

## Conclusion and Discussion

We have developed the R package *SpatialQPFs*, tailored for extracting features deciphering cell-cell spatial relationships from preprocessed digital pathology data. To the best of our knowledge, this is the first R package that comprehensively implements various spatial statistics methods and derives features based on cell coordinates and cell identities, comprehensively from point process data, areal data, and geostatistical data holistic perspective.

To validate its capabilities, we employed *SpatialQPFs* to extract features that quantify spatial interactions between tumor cells and lymphocytes in two distinct clinical trials. Subsequently, we investigated its capacity to predict patients’ survival risks, thereby showcasing its clinical relevance in discerning sub-populations that might derive greater benefits from cancer immunotherapy. This case study not only demonstrates the efficacy of *SpatialQPFs* in extracting meaningful insights from spatial data but also highlights its potential as a valuable tool in clinical research.

It’s worth noting that the functionalities of *SpatialQPFs* extend beyond analyzing H&E-stained digital pathology images to include other tissue imaging modalities such as Immunohistochemistry (IHC) and Immunofluorescence (IF) staining. For example, Li, X. et al.^59^ demonstrated this utility by analyzing IHC images. In their study, they used the *SpatialQPFs* package to derive spatial features that characterize the spatial distribution and colocalization of CD8 and pan-cytokeratin (CK) in lung and breast cancer cohorts, achieving the best performance for predicting immunophenotypes compared to other methods, including the state-of-the-art attention-based multiple instance learning (MIL)^60^. Another important application of *SpatialQPFs* is in the analysis of spatially resolved single-cell omics data. For instance, in a recent study by Ospina, O. et al.^18^, the authors developed an R package that leverages spatially-resolved gene expression data to provide spatial statistics, such as Moran’s I, Geary’s C, and Getis-Ord, to quantify tissue microenvironment heterogeneity. This approach captures transcriptomic complexity and enables the association with clinical outcomes. In another effort, after identifying cell types using -omics measurements, the authors^61^ used colocation quotient^62^ method which can be further improved by leveraging *SpatialQPFs* to investigate cell

type interactions, and to inform transcriptomics analysis using “avoidant” or “attractive” spatial relationships within the tissue microenvironment. These examples illustrate the broader utility of *SpatialQPFs* across different tissue imaging modalities and spatial data types.

Several limitations are worth noting. Not all researchers or individuals are proficient in R. With the development of artificial intelligence, Python^63^ has become arguably the most popular programming language. To broaden the adoption of the *SpatialQPFs* package, our strategy involves creating additional tutorials and vignettes accessible through Python wrapper functions. One promising solution for this endeavor might be *rpy*2, which provides an interface between R and Python. Future work will involve leveraging *rpy*2 to expand the accessibility of the package to a wider audience. Furthermore, since image preprocessing serves as prerequisite steps before invoking *SpatialQPFs*, it is essential to ensure that these preparatory procedures achieve satisfactory performance. For example, the performance of cell segmentation and classification must meet certain standards.

We believe that by leveraging computational tools like *SpatialQPFs*, researchers can effectively harness the vast potential embedded within pathology images, transforming them into a comprehensive array of human-interpretable and mineable variables on a large scale. This not only streamlines the process of analyzing vast amounts of data but also enables the identification of intricate patterns hidden within the tissue microenvironment. Consequently, this approach accelerates the discovery of novel imaging biomarkers relevant to understanding and diagnosing various human diseases. Through the systematic analysis facilitated by *SpatialQPFs*, researchers gain deeper insights into the complex interplay between cellular structures, paving the way for advancements in disease detection, prognosis, and treatment strategies.

## Acknowledgements

The author expresses gratitude to all members of Analytics and Medical Imaging, Product Development, Genentech, Inc. / F. Hoffmann-La Roche AG, and Computational Science and Informatics, Pathology Lab, Roche Diagnostic Solution for their invaluable feedback and assistance in curating the clinical images and outcome data. The clinical trials received support from Genentech, Inc. / F. Hoffmann-La Roche AG. Additionally, the author extends heartfelt thanks to the patients and their families.

## Author contributions statement

XL conceived and designed the study, conducted the experiments, and analysed the results.

## Additional information

### Supplementary Information

The Supplementary material is available at online version of the journal.

